# Binding of *Leishmania infantum* LPG to the midgut is not sufficient to define vector competence in *Lutzomyia longipalpis* sand flies

**DOI:** 10.1101/2020.06.10.145342

**Authors:** Iliano V. Coutinho-Abreu, James Oristian, Waldionê de Castro, Timothy R. Wilson, Claudio Meneses, Rodrigo P. Soares, Valéria M. Borges, Albert Descoteaux, Shaden Kamhawi, Jesus G. Valenzuela

**Affiliations:** Vector Molecular Biology Section, Laboratory of Malaria and Vector Research, National Institute of Allergy and Infectious Diseases, National Institutes of Health, Rockville, MD, USA; Fundação Oswaldo Cruz - FIOCRUZ, Instituto René Rachou, Belo Horizonte, MG, Brazil; Fundação Oswaldo Cruz - FIOCRUZ, Instituto Gonçalo Moniz, Salvador, BA, Brazil; INRS - Centre Armand-Frappier Santé Biotechnologie, 531 boul. des Prairies, Laval, Quebec, H7V 1B7, Canada

## Abstract

The major surface lipophosphoglycan (LPG) of *Leishmania* parasites is critical to vector competence in restrictive sand fly vectors by mediating *Leishmania* attachment to the midgut epithelium, considered essential to parasite survival and development. However, the relevance of LPG for sand flies that harbor multiple species of *Leishmania* remains elusive. We tested binding of *Leishmania infantum* wild type (WT), LPG-defective (Δ*lpg1* mutants) and add-back lines (Δ*lpg1* + *LPG1*) to sand fly midguts *in vitro* and their survival in *Lutzomyia longipalpis* sand flies *in vivo. Le. infantum* WT parasites attached to the *Lu. longipalpis* midgut *in vitro* with late-stage parasites binding to midguts in significantly higher numbers compared to early-stage stage promastigotes. Δ*lpg1* mutants did not bind to *Lu. longipalpis* midguts, and this was rescued in the Δ*lpg1* + *LPG1* lines, indicating that midgut binding is mediated by LPG. When *Lu. longipalpis* sand flies were infected with either *Le. infantum* WT, Δ*lpg1*, or Δ*lpg1* + *LPG1* of the BH46 or BA262 strains, the BH46 Δ*lpg1* mutant, but not the BA262 Δ*lpg1* mutant, survived and grew to similar numbers as the WT and Δ*lpg1* + *LPG1* lines. Exposure of BH46 and BA262 Δ*lpg1* mutants to blood engorged midgut extracts led to the mortality of the BA262 Δ*lpg1* but not the BH46 Δ*lpg1* parasites. These findings suggest that *Le. infantum* LPG protects parasites on a strain-specific basis early in infection, likely against toxic components of blood digestion, however, it is not necessary to prevent *Le. infantum* evacuation along with the feces in the permissive vector *Lu. longipalpis*.

**IMPORTANCE:** It is well established that LPG is sufficient to define the vector competence of restrictive sand fly vectors to *Leishmania* parasites. However, the permissiveness of other sand flies to multiple *Leishmania* species suggests that other factors might define vector competence for these vectors. In this study, we investigated the underpinnings of *Leishmania infantum* survival and development in its natural vector *Lutzomyia longipalpis*. We found out that LPG-mediated midgut binding persists in late-stage parasites. This observation is paradigm-changing and suggests that only a subset of infective metacyclics lose their ability to attach to the midgut with implications for parasite transmission dynamics. However, our data also demonstrate that LPG is not a determining factor in *Leishmania infantum* retention in the midgut of *Lutzomyia longipalpis*, a permissive vector. Rather, LPG appears to be more important in protecting some parasite strains from the toxic environment generated during blood meal digestion in the insect gut. Thus, the relevance of LPG in parasite development in permissive vectors appears to be complex and should be investigated on a strain-specific basis.

## Introduction

Phlebotomine sand flies (Diptera – Psychodidae) are biological vectors of *Leishmania* parasites (Kinetoplastidae). Different species of *Leishmania* cause a spectrum of diseases collectively known as leishmaniasis. Leishmaniasis is endemic in over 88 countries, putting over 350 million people at risk of becoming infected (1). Overall, 2 million people get infected with *Leishmania* annually, resulting in between 40,000 to 90,000 patient deaths due to complications from the most dangerous form of the disease, visceral leishmaniasis (1).

In order to establish a mature transmissible infection, *Leishmania* needs to escape from the barriers imposed by the sand fly midgut (2-8). Digestive enzymes in the sand fly gut were shown to be detrimental to the transitional stages during differentiation of *Leishmania major* amastigotes to procyclic promastigotes (5, 7). This susceptibility was also attributed to toxic byproducts from the digested blood for *Leishmania donovani* (9). Upon activation, the sand fly immune system is also known to be effective at reducing parasite loads by means of the Imd pathway (8). The peritrophic matrix (PM) represents another strong barrier to *Leishmania* development in the sand fly midgut (2, 3, 5); *Leishmania* relies on breakdown of the PM, mediated by a sand fly secreted chitinase, to escape its confinement (2). After this step, the parasites attach to the midgut epithelium (6, 10), and such attachment requires specific carbohydrate side chains on the surface of the parasite that bind a specific receptor on the midgut microvilli (6, 11).

Amongst the barriers preventing *Leishmania* development within the sand fly, midgut attachment has been suggested as the defining factor of sand fly vector competence (6, 11). The *Leishmania* surface is decorated with a rich glycocalyx (12-14), exhibiting four major types of GPI (glycosylphosphatidylinositol)-anchored glycoconjugates (15). The lipophosphoglycan (LPG) molecule is the most abundant component of the promastigote surface coat and consists of an oligosaccharide cap, a backbone of galactose-mannose repeated units (Gal(β1,3)Man(α1)-PO_4_), a conserved glycan core (Gal(α1,6)Gal(α1,3)Gal_f_(β1,3)[Glc(α1)-PO_4_]Man(α1,3)Man(α1,4)-GlcN(α1)), and a PI (1-*O*-alkyl-2-lyso-phosphatidylinositol)-anchor (13, 14, 16). The galactose-mannose repeats exhibit different carbohydrate side chains, depending upon the *Leishmania* species, strains, and stages (15, 17). It has been demonstrated that variations in the nature of side chain sugars decorating the LPG molecule of non-metacyclic stages confer specificity to interactions with vectors (15). Further, these side chains are modified to ensure release of metacyclics from the midgut (15); in some instances, the LPG molecule conformationally prevents binding of sugars to the midgut (15, 18-20). Such studies of vector competence, mostly undertaken under laboratory conditions, currently sorted sand flies into two groups (15). The restrictive, or specific, vectors, such *as P. papatasi, P. duboscqi* and *P. sergenti*, are able to support the development of only one species of *Leishmania* (15). The permissive sand fly vectors, such as *Phlebotomus perniciosus, Phlebotomus argentipes*, and *Lutzomyia longipalpis*, are capable of supporting multiple *Leishmania* species (21-23).

The functional binding properties of *Le. major* LPG have been extensively studied (6, 11, 24, 25). It has been demonstrated that the purified non-lipidic portion of the *Le. major* LPG (PG) binds only to midgut receptors of its natural sand fly vector, *Phlebotomus papatasi*, but binding is restricted to the LPG from the procyclic stage (6). Whereas the PG of the non-binding metacyclic stage is a much longer molecule and displays arabinose side chains (6), the procyclic LPG is shorter and exhibits galactose side chains (6), which are recognized by a galactose-binding lectin (PpGalec) expressed on the *P. papatasi* midgut epithelium (10). Supporting the importance of LPG in sand fly vector competence, a *Le. major* strain devoid of LPG side chains was not capable of developing in *P. papatasi* (11), nor was wild type *Le. major* fed along with anti-PpGalec antibodies (10). In both cases, the infections were mostly lost once the digested blood was passed. *Le. major lpg1* knockout parasites also failed to develop in another natural vector of *Le. major, Phlebotomus duboscqi* (24, 25).

Although the factors defining vector competence of the sand fly *P. papatasi* for *Le. major* are well known (6), they may not govern interactions of other sand fly-*Leishmania* pairs (11, 23-28). In fact, purified LPGs from multiple *Leishmania* species bind to the midgut of the sand fly *P. argentipes* (11), a permissive vector, despite exhibiting divergent carbohydrate side chains (15). Interestingly, *Le. major* and *Le. tropica* outcompete *Le. infantum* binding to midgut of its natural vector *Lu. longipalpis* when simultaneously in contact with the epithelium (28). In addition, the *Le. major lpg1* knockout line can successfully develop in both permissive *Lu. longipalpis* (24) and *P. perniciosus* (25) sand flies. Even though an LPG-independent mechanism based on a potential *Leishmania* lectin attaching to the microvilli glycocalyx has been proposed for *Leishmania* development in permissive vectors (23, 27), neither ligand nor receptor have been identified to date (29). Further, there is also a possibility that this phenomenon is restricted to *Le. major* development in permissive vectors and is not extendable to other *Leishmania* species naturally transmitted by such vectors. Such species-specific features have been demonstrated for the flagellar protein FLAG1/SMP1 that mediates midgut attachment of *Le. major* to *P. papatasi* but not *Le. infantum* to *Lu. longipalpis* (30, 31), and is further supported by the reduced survival of both *Le. infantum* and *Le. mexicana* Δ*lpg1* lines in *P. perniciosus* and *Lu. longipalpis*, respectively (26).

The mechanisms defining vector competence in permissive sand fly vectors is still an open question. The importance of LPG for midgut binding and parasite survival needs to be further analyzed, particularly as the lack of Δ*lpg1* + *LPG1* add-back lines in a previous study (26) precluded a definitive conclusion about the importance of LPG in vector competence of permissive vectors. Here, we assessed midgut binding, survival and development of *Le. infantum* in the sand fly *Lu. longipalpis* using multiple wild type strains as well as two Δ*lpg1* mutants and Δ*lpg1*+ *LPG1* add-back lines to answer the following questions: (1) Does *Le. infantum* bind to the midgut of *Lu. longipalpis*? (2) If it does, is the *Leishmania*-binding to the midgut stage-specific? (3) Is *Le. infantum* LPG necessary for parasite binding to the midgut? (4) Is the *Le. infantum* LPG sufficient to define vector competence in the natural permissive vector? (5) Do components of the blood bolus affect *Le. infantum* survival?

## Results

### *Le. infantum* binds to the midgut epithelium of *Lu. longipalpis*

In order to confirm previous results (6) and standardize the technique, we exposed *P. papatasi* midguts to *Le. major* (WR 2885) harvested from a 3-day-old culture. As expected, *Le. major* parasite bound to *P. papatasi* midguts (median 7,500 parasites/midgut; Fig 1A). Unexpectedly, *Le. infantum* LLM-320 (MHOM/ES/92/LLM-320) harvested on day 3 failed to bind to the midguts of the sand fly *Lu. longipalpis* (median 2,700 parasites/midgut; Fig. 1B). To our surprise, *Le. infantum* parasites from an older day 4 culture showed a greater number of parasites binding to *Lu. longipalpis* open midguts (median 17,850 parasites/midgut; Fig. 1C). As a control, 4 days old parasites did not bind intact, unopened, *Lu. longipalpis* midguts (median 1,600 parasites/midgut), pointing to the specificity of parasite binding to the sand fly midgut epithelium (Fig. 1C).

**Figure 1.**
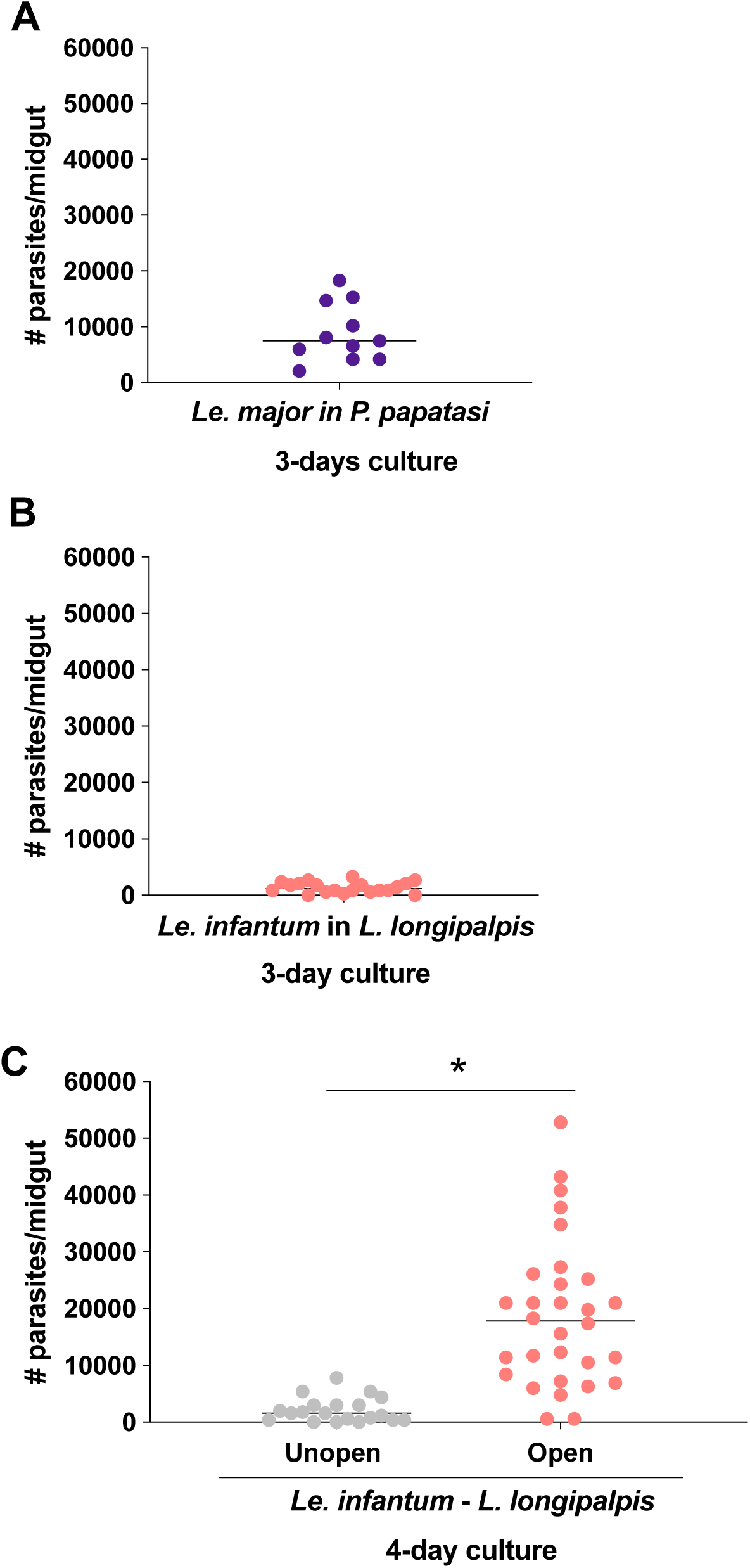
In vitro binding of *Leishmania major* and *Leishmania infantum* to midgut epithelia of natural sand fly vectors. (A-B) binding of *Le. major* (A) and *Le. infantum* LLM-320 (B) parasites, harvested from 3 days old cultures, to open midguts *Pheblotomus papatasi* and *Lutzomyia longipalpis* sand flies, respectively. n = 2. (C) Binding of *Le. infantum* parasites from cultures harvested at day 4 to unopened and opened *Lu. longipalpis* midguts. n = 2. Unopen midguts were intact; open midguts were cut longitudinally along the anterior-posterior axis. Unfed midguts were used. *, statistically significant at p < 0.05.

### *Le. infantum* binding to the midgut epithelium of *Lu. longipalpis* is stage-dependent

To validate and extend our observation, we used a different strain of *Le. infantum*, MCAN/BR/09/52 (LAB), and assessed its binding to sand fly midguts. Parasites harvested on day 3 attached to the midgut epithelium of *Lu. longipalpis* at a median of 5,250 parasites/midgut (Fig. 2A). As the LAB parasite cultures aged, the number of parasites binding the sand fly midgut increased proportionally, peaking with day 6 parasites at a median of 8,500 parasites/midgut (Fig. 2A). A greater proportion of parasite binding to the midgut epithelium as the culture aged was also observed for the *Le. infantum* BH46 strain (Fig. S1A-D), ranging from a median of 4,200 parasites/midgut for day 4 old culture parasites (Fig. S1A) to a median of 26,000 parasites/midgut with day 6 old culture parasites (Fig. S1C). As the culture got older (Fig. 2B; Fig. S2A), the parasites began to differentiate to the infective form, the metacyclic promastigotes, increasing from 10-30% on day 4 to about 80% on day 7 of culture, respectively (Fig. 2C; Fig. S2B). As we have observed that many, if not most, of the bound parasites were at late stages of differentiation (leptomonads and metacyclic), we used a Ficoll gradient to separate early-stage parasites (ESP) from late-stage parasites (LSP) and tested if they could bind to sand fly midgut epithelium. Surprisingly, we observed that significantly more LSP bound to the sand fly midgut epithelium compared to ESP, for both *Le. infantum* LAB (Fig. 2D and E) and BH46 (Fig. 2F and G) strains, harvested on both day 4 and day 5.

**Figure 2.**
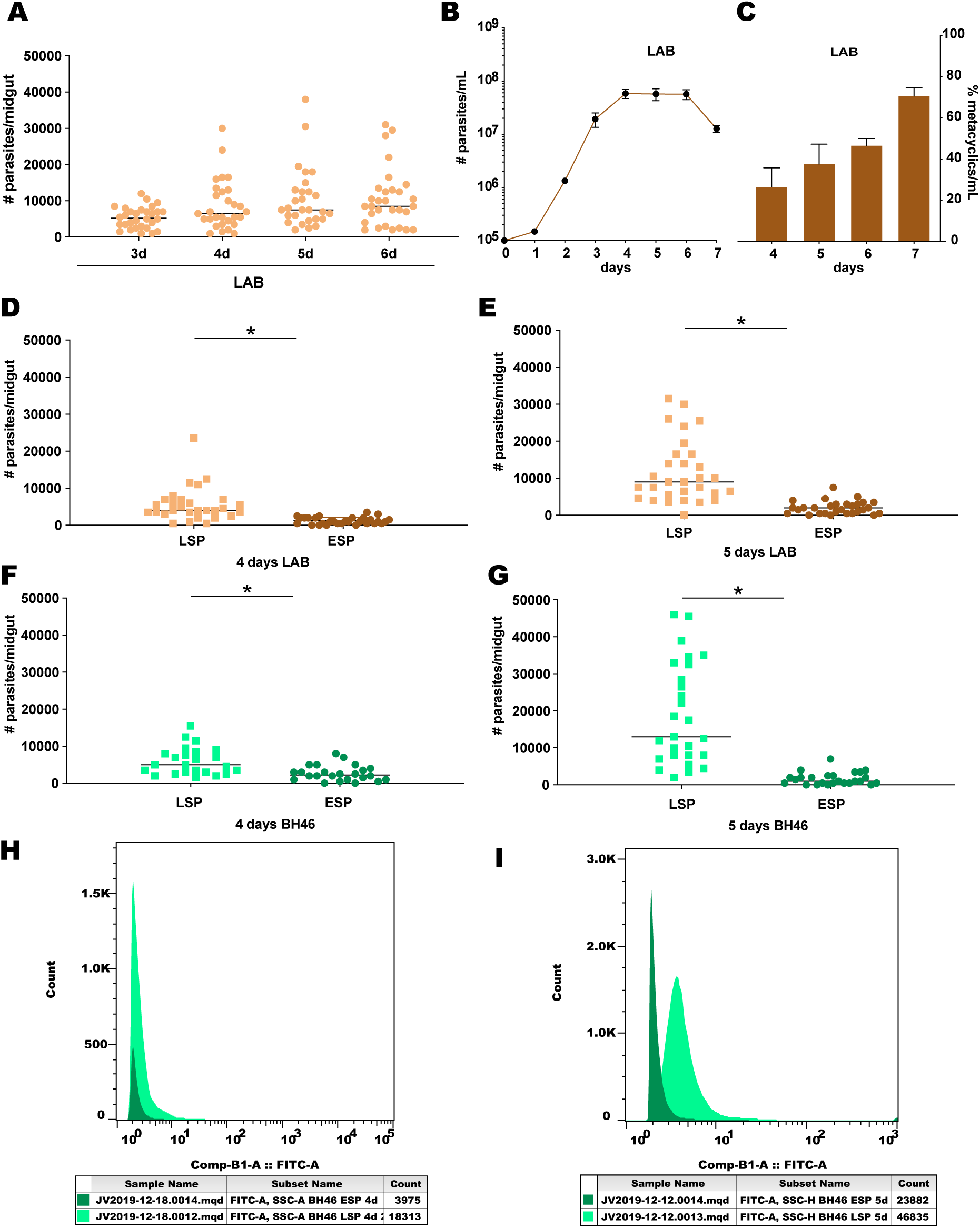
Temporal and stage specific binding of *Leishmania infantum* parasites to unfed *Lutzomyia longipalpis* midguts. (A-E) *Le. infantum* (MCAN/BR/09/52, LAB). (A) Binding of parasites harvested on different days of culture. n = 2. (B) Growth curve in culture. n = 3. (C) Metacyclic emergence on days 4-7 of culture. n = 3. (D-E) Differential binding of ESP and LSP from cultures harvested on days 4 (D) and 5 (E). n = 2. (F-G) Differential binding of ESP and LSP of *Le. infantum* BH46 wild type strain from cultures harvested on days 4 (F) and 5 (G). n = 2. ESP, early stage parasites; LSP, late stage parasites. Midguts were opened longitudinally along the anterior-posterior axis. *, statistically significant at p < 0.05. (H-I) Number of BH46 wild type parasites stained by anti-LPG antibody in metacyclic- (light green) and procyclic-enriched samples (dark green) for the 4 days (H) and 5 days (I) old parasite cultures.

In order to assess the abundance of LPG on the surface of BH46 WT parasites, we stained LSP and ESP Ficoll-purified parasites with a LPG backbone-specific monoclonal antibody (CA7AE). Using flow cytometry, we show that LSP parasites exhibited a 2- to 20-fold higher antibody binding than ESP samples for 4 and 5-day old cultures, respectively (Fig. 2H-I), indicative of the increased abundance of LPG in the former. The increased fluorescence intensity as the parasites aged (Fig. 2H-I) correlated with a greater number of parasites binding the sand fly midgut from 4d and 5d old cultures (Fig. 2F-G). In order to further evaluate such observations, we measured LPG abundance in these parasites by confocal microscopy (Fig. 3A-H). We observed a more intense LPG staining in LSP (Fig. 3A-D) compared to ESP (Fig. 3E-H). In comparison, 3-day old parasite cultures stained poorly with the CA7AE antibody (Fig. 3I-L).

**Figure 3.**
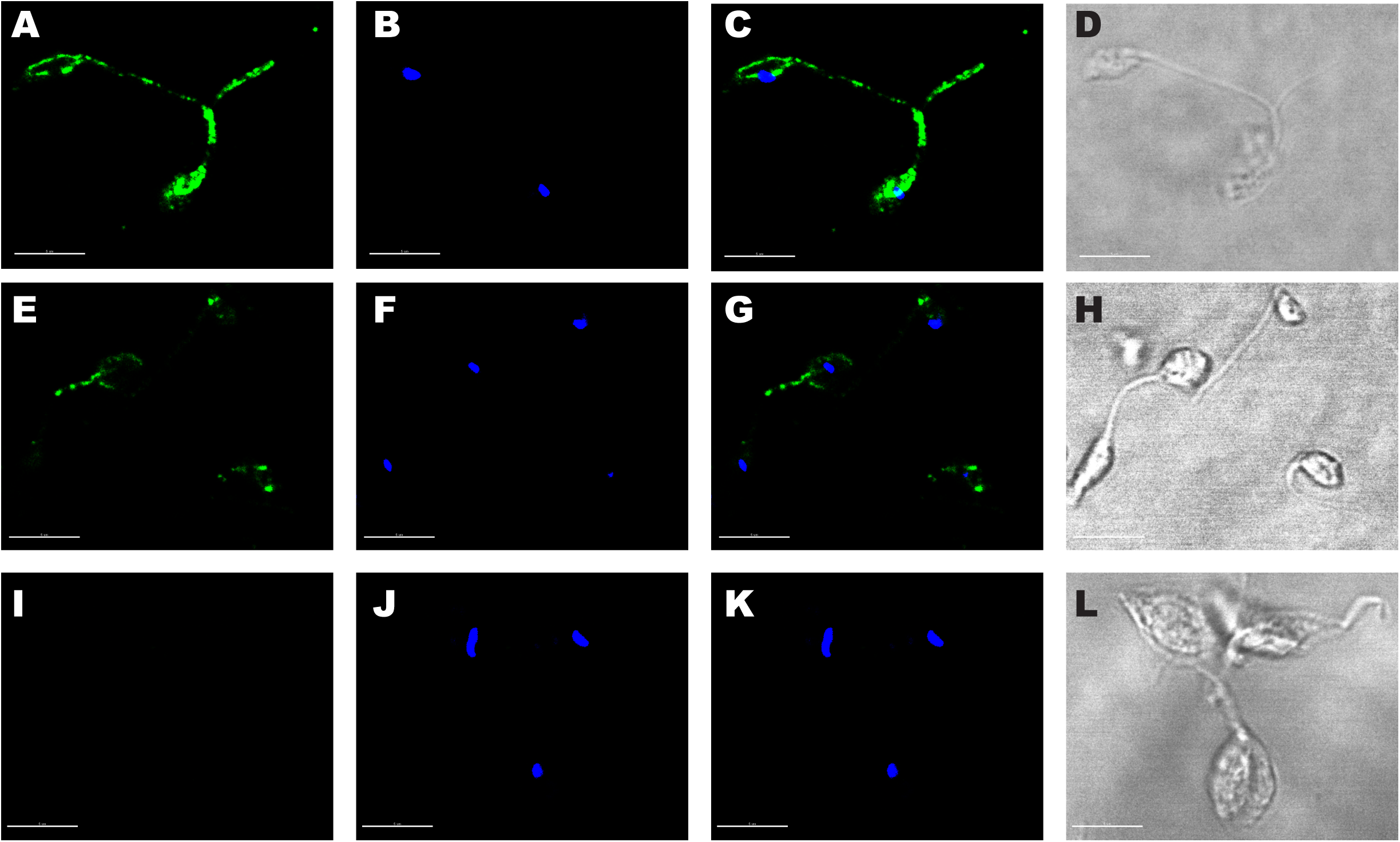
Immunostaining of LPG on the surface of different *Leishmania infantum* BH46 developmental stages. (A-L) Wild type parasites were stained with GFP-conjugated anti-LPG CA7AE antibody. (A-H) Seven days old cultures were harvested, and parasites were sorted into metacyclic (A-D) and procyclic (E-H) promastigotes in a Ficoll gradient. (I-L) Staining of three days old culture parasites. Bar = 5 µm. Green = LPG. Blue = DAPI (nuclear DNA). Gray pictures = DIC.

### *Le. infantum* binding to the midgut epithelium of *Lu. longipalpis* is LPG-dependent

In order to assess whether or not the major surface glycoconjugate of *Le. infantum* (LPG) was the parasite ligand to the midgut epithelium of *Lu. longipalpis*, we carried out midgut binding assays with the *Le. infantum* BH46 wild type strain (BH46 WT) as well as both the LPG-deficient (Δ*lpg1*) mutant and the add-back (Δ*lpg1* + *LPG1*) lines. Similar to what was observed for unfed midguts (Fig. S1), aging parasites from the BH46 WT line bound to the epithelium of midguts dissected 5 days after blood feeding with increasing efficiency, whereas parasite binding was limited in unopened midguts (Fig. 4). For both fed and unfed *Lu. longipalpis* midguts, binding of the BH46 Δ*lpg1* + *LPG1* was intermediate and significantly higher than that of the BH46 Δ*lpg1* mutant that failed to bind, exhibiting less than 1,500 parasite/midgut regardless the age of the harvested parasites or the feeding status of the midguts (Fig. 4 and Fig. S1). Similar results were obtained with the *Le. infantum* BA262 Δ*lpg1* mutant, which also failed to bind to unfed midguts (Fig. S3). Comparatively, the BA262 WT bound with increased efficiency as the parasites aged (Fig. S3 and S4). Despite the low midgut binding of the BA262 Δ*lpg1* + *LPG1*, such parasites bound to midguts at significantly greater proportions than the BA262 Δ*lpg1* mutant (Fig. S3).

**Figure 4.**
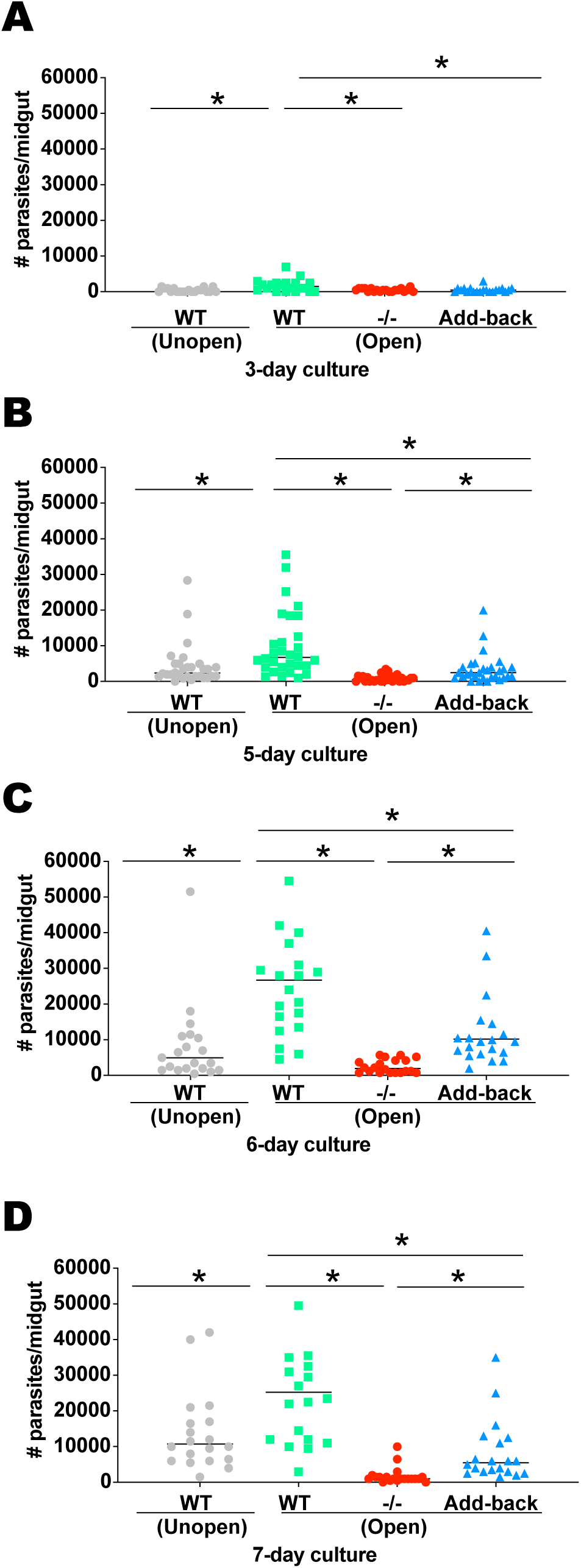
Binding of *Leishmania infantum* BH46 wild type, Δlpg1, and Δlpg1 + LPG1 lines to *Lutzomyia longipalpis* midguts dissected 5 days after blood feeding. (A-D). Cultures of *Le. infantum* BH46 WT (wild type), -/- (Δlpg1), and Add-back Δlpg1 + LPG1) were harvested on days 3 (A), 5 (B), 6 (C), and 7 (D) and incubated with midguts of sand flies 5 days after blood feeding. Midguts were opened longitudinally along the anterior-posterior axis. WT strain was also incubated with intact (unopened) midguts. n = 2. *, statistically significant at p < 0.05.

### *Le. infantum* BH46 Δ*lpg1* mutants grow in the midguts of *Lu. longipalpis*

As *Le. infantum* LPG is sufficient to mediate midgut epithelium attachment *in vitro*, but binding was substantially increased in LSP compared to ESP, we tested whether this ligand is necessary for *in vivo* parasite development in the midgut of the sand fly *Lu. longipalpis*. We infected *Lu. longipalpis* sand flies with the BH46 WT, BH46 Δ*lpg1*, and BH46 Δ*lpg1* + *LPG1* and followed both parasite load and infection prevalence at five time points after infection. When seeded with 5 million parasites/mL, the BH46 Δ*lpg1* mutant exhibited a significantly lower parasite load and a lower infection prevalence compared to BH46 WT strain or to the BH46 Δ*lpg1* + *LPG1* on day 3 post infection (Fig. 5A-B), but it recovered on subsequent days, exhibiting a similar parasite load and infection prevalence to the ones infected with either BH46 WT or BH46 Δ*lpg1* + *LPG1* (Fig. 5C-J). When the infection was started with 2 million parasites/mL, the BH46 Δ*lpg1* mutant displayed a lower, yet not significant, parasite load on day 3 (Fig. S5A), but higher parasite loads than either the WT strain or the Δ*lpg1* + *LPG1* from day 6 onwards (Fig. S5B-E).

**Figure 5.**
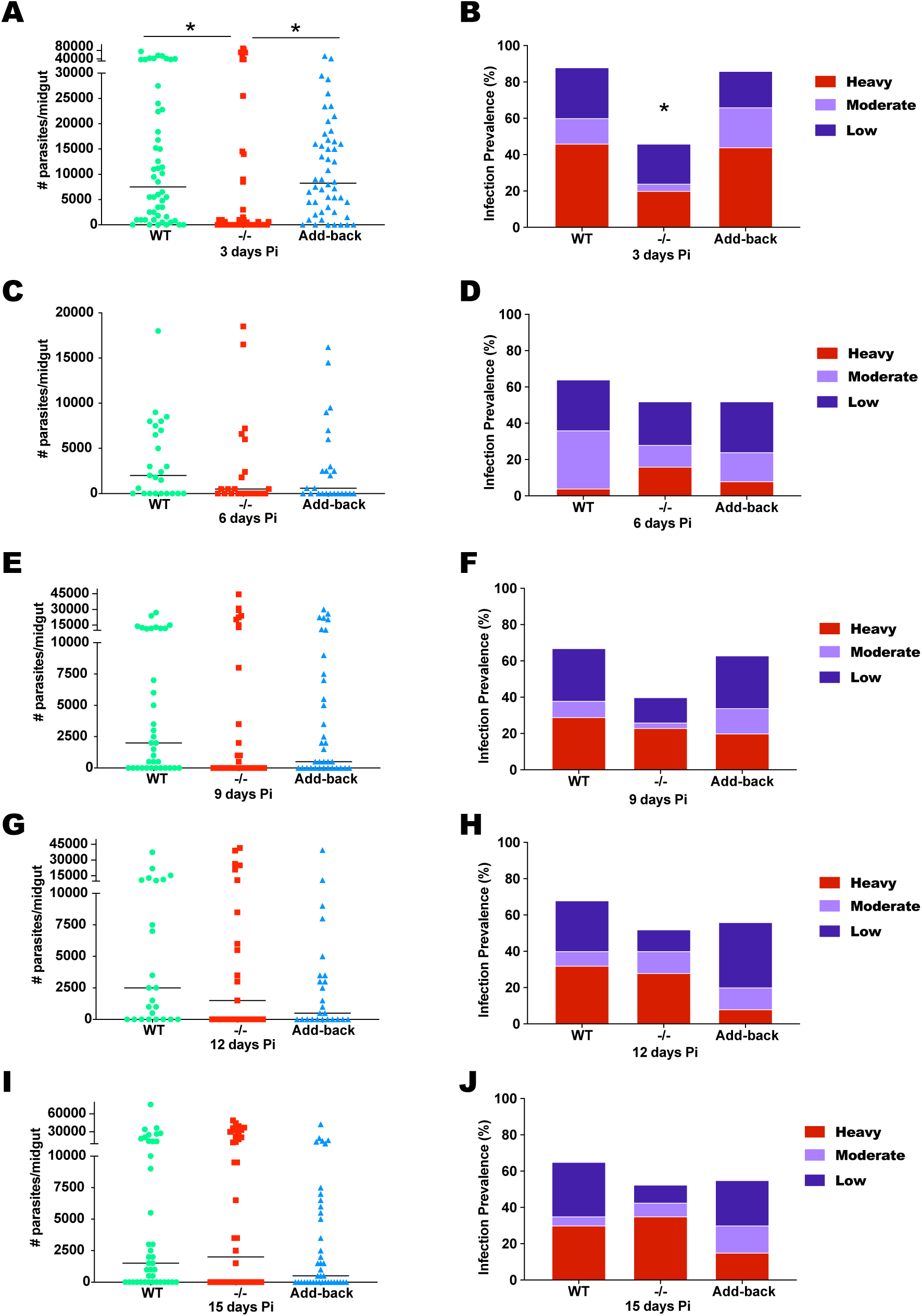
*Lutzomyia longipalpis* midgut infection with *Leishmania infantum* BH46 wild type, Δlpg1, and Δlpg1 + LPG1 parasites. (A-J) Upon infection with 5 million parasites per mL, parasite load and infection prevalence were assessed on days 3 (A-B), 6 (C-D), 9 (E-F), 12 (G-H), and 15 (I-J) after infection, respectively. WT (wild type), -/- (Δlpg1), and Add-back (Δlpg1 + LPG1). Low: 500-5,000 parasites/midgut; Moderate: 5,000-10,000 parasites/midgut; Heavy: > 10,000 parasites/midgut. n = 2. *, statistically significant at p < 0.05.

### *Le. infantum* BA262 Δ*lpg1* mutants fail to grow in the midguts of *Lu. longipalpis*

As the LPG side chain decorations are polymorphic amongst *Le. infantum* strains (32), we also assessed the ability of BA262 WT strain, along with the BA262 Δ*lpg1* and BA262 Δ*lpg1* + *LPG1* lines (33), to develop in *Lu. longipalpis* (Fig. 6). Different from the BH46 strain, which exhibits side chains with 1 to 3 β-glucose residues, the LPG of BA262 is devoid of side chains, like most of the *Le. infantum* strains (32). Seeded with 5 million parasites/mL, the BA262 Δ*lpg1* mutant displayed lower parasite loads on days 2 and 3 (median 0-500 parasites/midgut) as compared to the WT (median 3,000-5,000 parasites/midgut) and Δ*lpg1* + *LPG1* (median 1,600-13,750 parasites/midgut) lines, and a lower infection prevalence before the blood was passed (Fig. 6A-D). After the blood meal was passed, the BA262 Δ*lpg1* mutant was lost in the majority of sand flies, persisting in only a few specimens that displayed a reduced number of parasites and a decreased infection prevalence (Fig. 6E-L). The BA262 WT and Δ*lpg1* + *LPG1* lines, on the other hand, developed well in the midguts, reaching a median of 27,000 and 35,000 parasites per midgut, respectively, an 80 to 85% infection prevalence on day 15 post infection (Fig. 6K-L).

**Figure 6.**
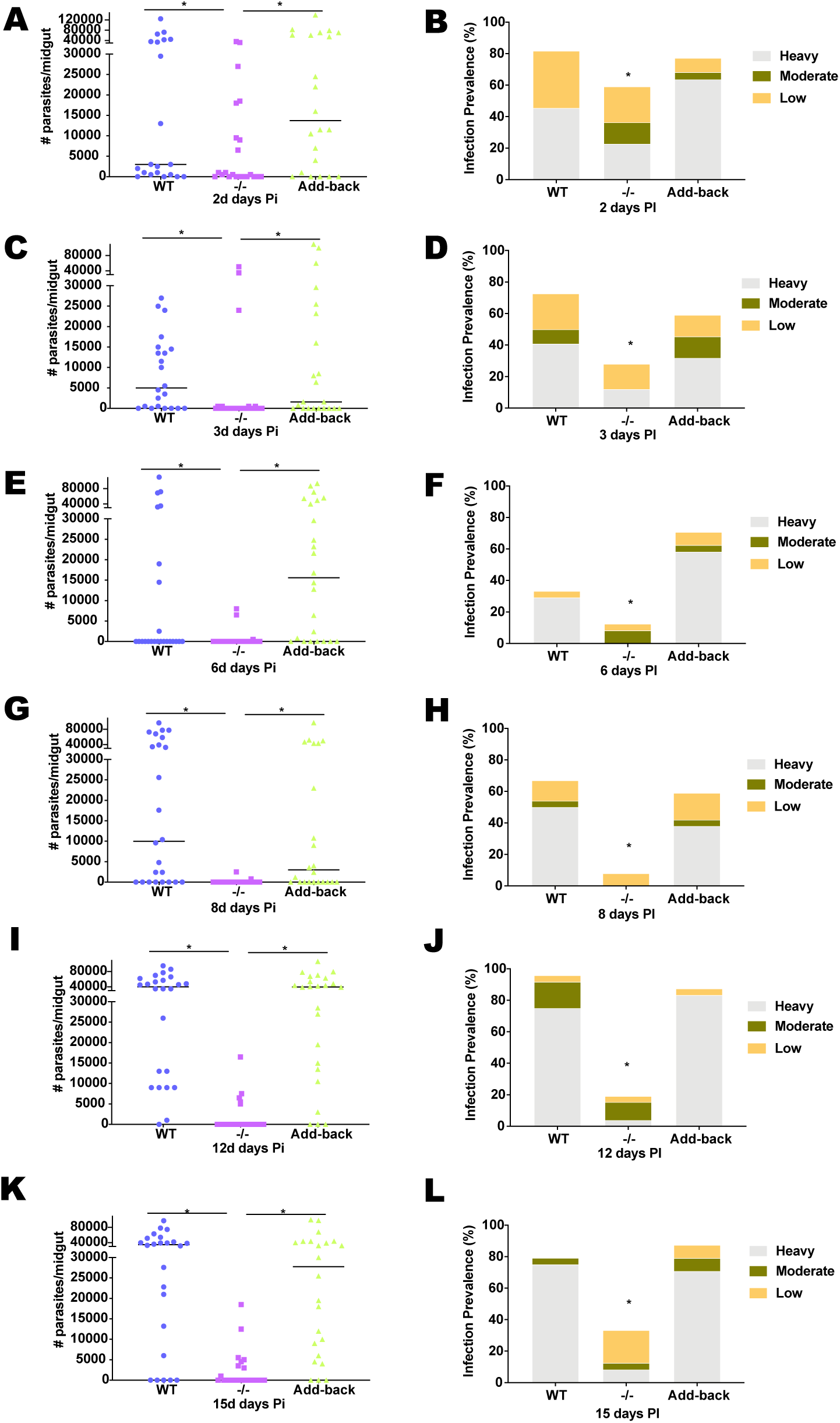
*Lutzomyia longipalpis* midgut infection with *Leishmania infantum* BA262 wild type, Δlpg1, and Δlpg1 + LPG1 parasites. (A-J) Upon infection with 5 million parasites per mL, parasite load and infection prevalence were assessed on days 2 (A-B), 3 (C-D), 6 (E-F), 8 (G-H), 12 (I-J), and 15 (K-L) after infection, respectively. WT (wild type), -/- (Δlpg1), and Add-back (Δlpg1 + LPG1). Low: 500-5,000 parasites/midgut; Moderate: 5,000-10,000 parasites/midgut; Heavy: > 10,000 parasites/midgut. n = 2. *, statistically significant at p < 0.05.

### *Le. infantum* BH46 and BA262 Δ*lpg1* mutants display different susceptibilities to components of the blood meal

To investigate the differences in sand fly survival of the BH46 and BA262 Δ*lpg1* mutants, we tested if resistance to by-products blood meal digestion could be a factor that explains these differences. For this, we incubated *in vitro* parasites in exponential phase of growth with extracts of midguts, collected at 24 and 48 hours after blood feeding, for 4 hours at 26°C, and compared them to WT and Δ*lpg1* + *LPG1* lines (Fig. 7). BH46 Δ*lpg1* mutants were not affected by the components of the digested sand fly blood meal (Fig. 7A). In contrast, the BA262 Δ*lpg1* mutants were severely affected by incubation with midguts collected 24h and 48h post-blood feeding (Fig. 7B). The WT and Δ*lpg1* + *LPG1* lines of both strains were not affected by incubation with midgut extracts (Fig. 7A-B).

**Figure 7.**
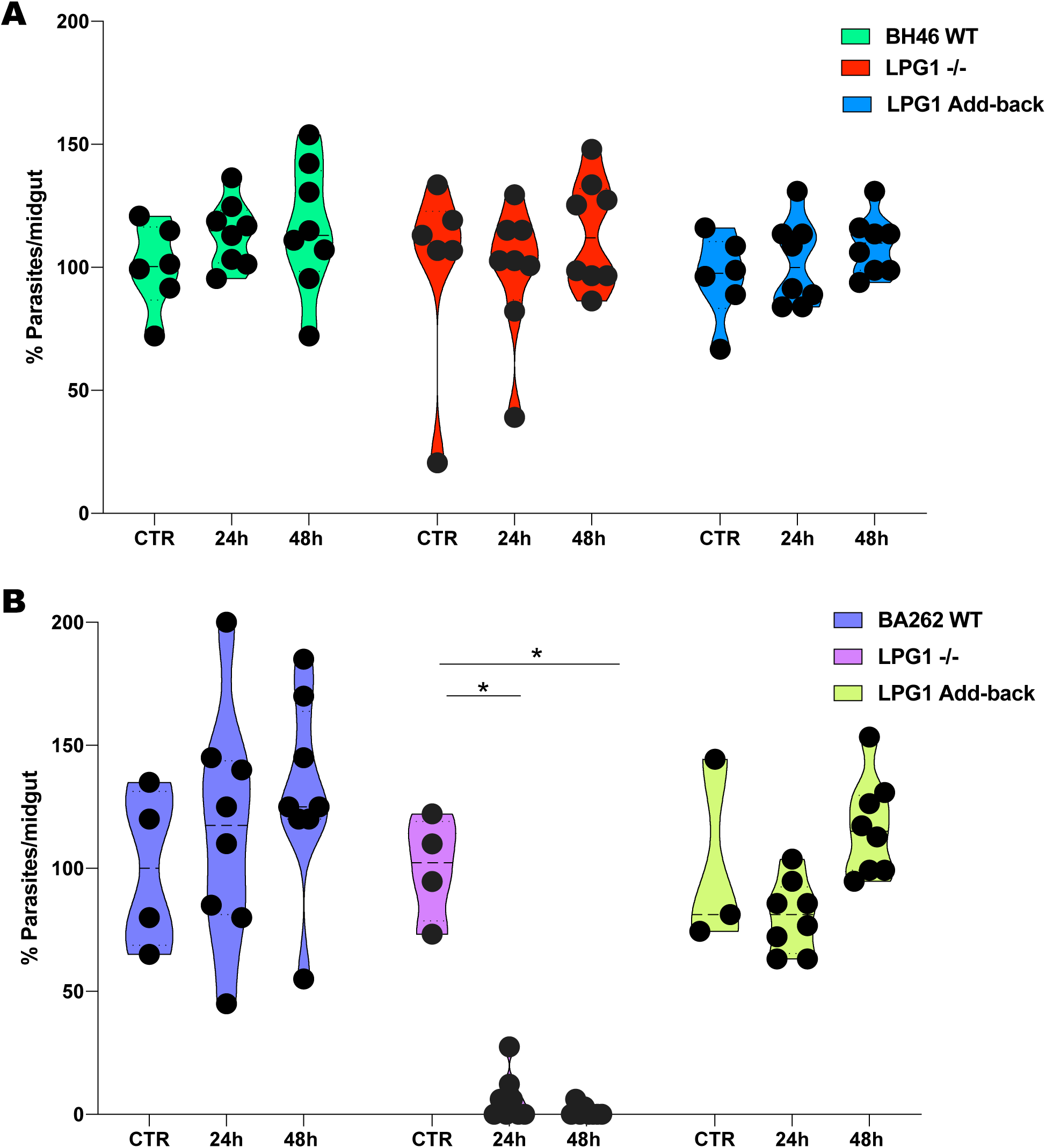
*Leishmania infantum* survival after incubation with extracts of blood engorged *Lutzomyia longipalpis* midguts in vitro. (A) BH46 WT (wild type), -/- (Δlpg1), and Add-back (Δlpg1 + LPG1) lines were exposed in vitro to either PBS (CTR) or extracts of engorged midguts dissected at 24h or 48h post blood meal. (B) BA262 WT (wild type), -/- (Δlpg1), and Add-back (Δlpg1 + LPG1) parasites were exposed in vitro to either PBS (CTR) or extracts of engorged midguts dissected at 24h or 48h post blood meal. n = 2. *, statistically significant at p < 0.05.

## Discussion

Many studies have demonstrated that binding by *Le. major* to the midguts of the restrictive vectors is mediated by LPG (6, 10, 11, 23-25, 34). Incubation of *P. papatasi* midguts with purified PGs from procyclic *Le. major* parasites prevented the binding of procyclic *Le. major* parasites (6), and *Le. major* Δ*lpg1* parasites cannot develop in *P. papatasi* (23, 34) or *P. duboscqi* (24, 25) sand flies. On the other hand, the *Le. major* Δ*lpg1* binds to midguts and develops well in permissive vectors, such as *Lu. longipalpis* (23, 24), as well as *Phlebotomus arabicus* (23), *P. argentipes* (25), and *P. perniciosus* (25). Based on such findings, an LPG-independent mechanism for midgut binding was proposed for the permissive vectors (27). Differences in midgut glycosylation between restrictive and permissive vectors were observed that could possibly account for non-specific binding of parasites in the latter. It was hypothesized that a lectin on the *Leishmania* surface might bind to the *O*-linked glycans of the midgut microvilli of permissive vectors allowing parasite binding (25, 27). Nonetheless, such studies were carried out using *Le. major* and unnatural permissive vectors (23-25, 27); thereby, the LPG-independent mechanism could be restricted to *Le. major* development in unnatural permissive vectors. Therefore, we decided to revisit the role of LPG in parasite development in natural permissive vectors, focusing on two *Le. infantum* strains bearing intraspecies polymorphisms in their LPGs (25). The LPG of one strain, BA262, is devoid of side chains (type I) whereas that of the BH46 strain has β-glucose sugars branching-off the repeat unit backbone (type III; (20)). Here, we demonstrate that *Le. infantum* (BH46) wild type and Δ*lpg1* + *LPG1* lines bound to both unfed and 5 days post-feeding midgut epithelia of *Lu. longipalpis in vitro*. In contrast, the LPG-deficient Δ*lpg1* failed to bind to *Lu. longipalpis* unfed and fed midguts. Similar results were reported for Δ*lpg1* mutants of *Le. mexicana* (MPRO/BR/72/M1845) and *Le. infantum* (MHOM/BR/76/M4192) that exhibited poor binding to midguts of *Lu. longipalpis* and *P. perniciosus* (26), respectively. Altogether, these data indicate that LPG mediates *Leishmania* binding to midguts of natural permissive vectors, and suggest that the previously described LPG-independent midgut binding mechanism may be limited to *Le. major* binding to midguts of permissive vectors.

Similar to conclusions for *Le. major* binding to the midgut of its natural restrictive vector, previous reports claimed that *Le. infantum* and *Le. donovani* binding to the midgut of a natural permissive vector was also restricted to early-stage parasites (19, 20). Nonetheless, the fact that PNA was used to sort procyclic and metacyclic *Le. infantum* (20) and *Le. donovani* (19) parasites may have confounded interpretation of results. PNA has been used to purify *Le. major* metacyclics as the LPGs of such parasite stages are decorated with arabinose that replaces numerous β-galactose sugars on side chains of early stage parasites (35). In contrast, *Le. infantum* (20) and *Le. donovani* (19) LPGs only display a single β-galactose residue in the cap. Whereas it is not clear whether or not the same residue is absent in the *Le. infantum* metacyclic stage LPG cap (20), the *Le. donovani* cap bears β-galactose or mannose residues at the same proportions as procylic parasites (19), precluding the accuracy of procyclics and metacyclics purification by PNA. Additionally, experiments showing the developmental differences of *Le. infantum* (20) and *Le. donovani* LPG (19) found that PNA-purified metacyclic LPG was representative of only 10% of the parasites in stationary phase cultures (19, 20). It is possible that PNA-purified metacyclic LPGs are representative of a small metacyclic subpopulation that does not bind to PNA, as the whole metacyclic population usually accounts for about 80% of late phase cultures. Importantly, this small population of non-binding/free swimming *Leishmania* parasites likely comprise the metacyclics that are transmitted, as the parasites inoculated by sand flies represent only 0.1% to 14% of a mature *Leishmania* infection in sand flies (36). The observed increase in the intensity of LPG labelling as *Le. infantum* aged in culture, which correlated positively with the number of bound parasites to midguts, supports this hypothesis. Knowing whether stronger LPG labelling is related to an increase in the number and/or length of LPG molecules during metacyclogenesis will shed light on the binding mechanism of the *Le. infantum* LPG to the receptor on the midgut microvilli.

Electron microscopy images of *Le. infantum* developing in *Lu. longipalpis* show that parasites, then described as long and short nectomonads, and now termed nectomonads and leptomonads, respectively, were observed attached to the posterior and anterior midguts throughout the parasite life cycle in the sand fly (37). In our experiments, early-stage parasites were not able to attach to midguts *in vitro*, which might be explained by a lack of *bona fide* nectomonad parasites in cultures expressing LPG on their surface. On the other hand, we observed strong *Lu. longipalpis* midgut binding by late-stage *Le. infantum* parasites, enriched in leptomonads and metacyclics, confirming early observations by electron microscopy that *Le. infantum* late-stage parasites also bind to the *Lu. longipalpis* midgut epithelium (37). Whether midgut binding by late stage parasites is necessary for parasite migration to the anterior midgut (38), genetic exchange (39, 40), and/or preventing metacyclic parasites to be pushed to posterior midgut upon sequential blood meals (41), has yet to be determined.

Our proposed model of LPG-mediated *Le. infantum* attachment to the midgut of a permissive vector complements and expands upon previous findings demonstrating that changes in LPG during metacyclogenesis mediates parasite detachment from the midgut epithelium to be transmitted. Here, we propose that metacyclics encompass two subpopulations: one that binds to the midgut epithelium, as was observed in this study and also earlier (37), and one free-swimming that is transmitted by a sand fly bite (19, 20). The existence of such subpopulations is also supported by polymorphisms in the LPG cap of metacyclic *Le. infantum* (20) and *Le. donovani* (19) parasites.

It was previously shown that lack of LPG in *Le. infantum* seems to affect the ability of this parasite to develop in the midgut of the permissive natural vector *P. perniciosus* (26). A closer look at the results, nonetheless, suggests that the *Le. infantum* Δ*lpg1* infection shows some recovery on day 8 post infection, after the digested blood was passed, compared to day 4, when the blood meal is still being digested, as the prevalence of sand flies infected with heavy and moderate infections increased (26). In order to further investigate the importance of LPG for *Le. infantum* development in *Lu. longipalpis*, we assessed survival and development of the *Le. infantum* BH46 and BA262 WT, Δ*lpg1*, and Δ*lpg1* + *LPG1* lines in the natural permissive vector *Lu. longipalpis* for 15 days. In accordance with the previous report (26), the *Le. infantum* BH46 Δ*lpg1* mutant struggled to survive at the time of blood digestion on day 3, yet it thrived after the blood was passed, reaching similar parasite loads as the WT and add-back lines at later time points. In contrast, the BA262 Δ*lpg1* mutant not only struggled to survive up to day 3 but succumbed afterwards, revealing important inter-strain differences in *Le. infantum in vivo* development. These data indicate that LPG-mediated epithelium binding is not a determinant of parasite survival in the midgut for this parasite-vector pair. However, the presence of LPG does confer protection to some *Le. infantum* strains such as BA262.

Toxicity of blood components can affect the survival of *Leishmania* parasites in the sand fly midgut during digestion (9). In the present study, incubation of the *Le. infantum* strains, BH46 and BA262, harvested from 3 days old cultures with extracts of engorged midguts produced different outcomes. Whereas the BH46 and BA262 WT and Δ*lpg1* + *LPG1* lines, as well as the BH46 Δ*lpg1* mutant, survived exposure to the toxic components of the digested blood, the BA262 Δ*lpg1* mutant was highly affected by midgut extracts. These results correlated well with the survival of the BH46 Δ*lpg1* mutant and the lack of further development of the BA262 *lpg1* KO line in sand flies. Together, these results indicate that neither LPG type I nor III is needed to prevent *Le. infantum* from being eliminated along with the digested blood bolus in a permissive vector, but it is likely important for early survival during the digestive period for some but not all parasite strains. The survival of the BH46 Δ*lpg1* mutant after exposure to engorged midgut extracts is intriguing; the nature of the glycosylation of the galactose-mannose backbone of other surface glycoconjugates may provide protection for this strain and needs to be further explored.

As stated by Sacks and Kamhawi (15), for permissive species such as *Lu. longipalpis*, persistence of parasites in the midgut after blood is passed is possibly due to factors other than attachment, such as a slower peristaltic movement. These observations are supported by our findings which indicate that LPG-docking of parasites does occur but appears not to define vector competence for *Le. infantum* in *Lu. longipalpis*. Rather, vector competence in permissive vectors seems to be more complex, affected by strain-specific differences and involving multiple stages of parasite development in the midgut.

## Materials and Methods

### Sand flies, *Leishmania* strains, and parasite cultivation

The sand flies used in this study belonged to either the *Lu. longipalpis* Jacobina Colony or the *P. papatasi* Jordan colony, both maintained at the LMVR/NIH sand fly insectary. The different *Le. infantum* strains LAB (MCAN/BR/09/52, (42)), BH46 wild type (BH46 WT, MCAN/BR/89/BH46), the LPG-deficient BH46 Δ*lpg1* and the BH46 Δ*lpg1* + *LPG1* add-back (BH46 res), BA262 (MCAN/BR/89/BA262, (33)), the LPG-deficient BA262 Δ*lpg1* and the BA262 Δ*lpg1* + *LPG1* (BA262 res), RFP (MHOM/ES/92/LLM-320, RFP expressing (43)), *Le. major* (WR 2885, RFP-expressing; (36)) were cultivated in Grace medium (Lonza Biowhittaker and Sigma-Aldridge) supplemented with 20% heat-inactivated fetal calf serum (FCS; Gibco, 16140071) and pen/strep 100ug/mL in 25 cm^2^ flasks. For BA262, FCS was further heated inactivated at 56°C for 1 hour. For all the experiments, cultures were seeded with 1 × 10^5^ parasites/mL, and parasites were grown at 26°C in a B.O.D chamber. For both BH46 -/- and BH46 Add-back, Hygromycin (50ug/mL) and Geneticin (G418, 5ug/mL) were added to the medium. In addition, Zeocin (75ug/mL) was added to BH46 Add-back cultures. For both BA262 Δ*lpg1* and BA262 Δ*lpg1* + *LPG1*, Hygromycin (50ug/mL) and Geneticin (G418, 70ug/mL) were added to the medium. In addition, Zeocin (100ug/mL) was added to BA262 Δ*lpg1* + *LPG1* cultures.

### Sorting of early-stage and late-stage parasites in Ficoll gradient

Procedure was performed as described elsewhere (44), with slight modifications. Briefly, cultures were spun down, and parasites were washed twice in PBS and resuspended in 2mL PBS. Then, parasites were overlaid with 40% Ficoll in 2mL PBS, followed by addition of 10% Ficoll in 2mL M199 medium, and spun at 365 x g for 10 min at room temperature. Metacyclic-enriched parasites were collected from the layer in the interface between 10% Ficoll and PBS whereas procyclic-enriched parasites were collected after removing supernatant and resuspending the pellet. Thereafter, both parasite samples were washed twice in PBS for further experimentation.

### Parasite Growth Curves

Parasite cultures were seeded with 1 × 10^5^ parasites/mL as described above, and parasites were counted daily direct from the medium or diluted in PBS using Neubauer improved chambers (Incyto).

### Midgut binding assays

Parasite cultures were spun down once and washed with PBS twice at 3,500 RPMs for 15 minutes. Parasites were counted and diluted to 5 × 10^7^ parasites/mL in PBS. The midguts of *P. papatasi* and *Lu. longipalpis* unfed as well as *Lu. longipalpis* 5 days after blood feeding were dissected in a drop of PBS, opened up transversally in Y shape with fine entomological pins when needed, and groups of 10 midguts were exposed to 2 × 10^6^ parasites in 40uL of PBS in a well of an electron microscopy 9 cavity Pyrex pressed plate (Fisher Scientific). Chambers were transferred to a humidified chamber, and incubation was performed for 45 min at room temperature. Afterwards, midguts were passed through fresh PBS twice and transferred individually to 1.7mL Eppendorf tubes (Denville Scientific) with 30uL of PBS.

### Sand fly infections

Defibrinated naïve rabbit blood (Noble Life Sciences, Gaithersburg, MD), was spun down at 2,000 RPMs for 10 min, and plasma was collected and transferred to a fresh vial. RBCs were washed at least twice with PBS (until most of the free heme was removed) whereas plasma was heat-inactivated at 56°C for 1 hour. Parasite cultures were spun down and washed twice with PBS as described above. Then, RBCs were reconstituted with heat-inactivated plasma and seeded with either 5 × 10^6^ (BH46 and BA262) or 2 × 10^6^ parasites/mL (BH46). Infectious blood was loaded into a custom-made glass feeder (Chemglass Life Sciences, CG183570), capped with a chick skin and heated by a circulating water bath set for 37°C. Sand flies were allowed to feed for 3 hours. Afterwards, midguts were dissected and transferred individually to 1.7mL Eppendorf tubes (Denville Scientific) in 50uL PBS.

### Midgut *Leishmania* load assessment

Midguts were homogenized with a cordless motor and disposable pellet mixers (Kimble). In order to assess metacyclic proportions, formalin was added to the vials at a 0.005% final concentration. Samples were diluted as necessary and 10uL was loaded onto Neubauer improved chambers (Incyto).

### *Leishmania* incubation with extracts of blood engorged midguts

For sand fly feeding on naïve rabbit blood (Noble Life Sciences, Gaithersburg, MD), RBCs and plasma were processed as described above. Sand flies were fed on the naïve (heat-inactivated) blood, and midguts were dissected at 24 and 48 hours after feeding. Midguts were individually transferred to 0.2mL tubes, frozen in dry ice, and stored at −80°C. Two batches of midguts were obtained from sand flies fed on at two different days. Before incubation with parasites, midguts underwent 10 cycles of freeze-thaw (dry ice/room temperature; 5 minutes each). Parasites were harvested from 3 days old culture, washed twice in 1X PBS, and diluted to 5 × 10^6^ parasites/mL in complete Grace medium. One microliter (5,000 parasites) was incubated with either the extract of a single midgut or PBS (1ul) for 4 hours at 26°C. Afterwards, 20uL of PBS were added to each sample, and parasites were counted with Neubauer improved chambers.

### Confocal Microscopy

*Le. infantum* BH46 WT line was grown as above describe and harvested at days 3 and 7. For day 7 parasites, procyclics and metacyclics were isolated in a Ficoll gradient as described above. In μ-Slide Angiogenesis (Ibidi Cells in Focus, 81506) coated with Poly-L-lysine (Sigma Aldridge, P8920), one million parasites were fixed in 4% paraformaldehyde on ice for 15 minutes, blocked with goat serum (Sigma Aldridge, G9023) for 1 hour, incubated with the primary antibody CA7AE (1μg/μL) at 1:1,000 dilution for 30 minutes, and the secondary antibody Alexa Fluor® 488 goat anti-mouse IgG (H+L; Molecular Probes A11001) at 1:5,000 dilution for 30 minutes. Samples were mounted with DAPI containing Flouromont-G (Southern Biotech, 0100-20). Images were obtained with the Leica TSC SP5 in z-stacks of 0.42μM. Images were analyzed with the Imaris software (Oxford Instruments).

### Flow Cytometry

Metacyclic- and procyclic-enriched samples were sorted from both BH46 and BA262 4 and 5 days old cultures by ficoll gradient centrifugation as described above. One million parasite of each sample/strain were washed twice in cell staining buffer (BioLegend, 420201) and resuspended in 100 μL of the LPG backbone specific CA7AE antibody at 0.125μL/100μL dilution in staining buffer. Samples were incubated with primary antibody for 30 minutes at 4°C, washed twice with staining buffer, and then incubated with the secondary antibody Alexa Fluor® 488 goat anti-mouse IgG under the same conditions. After this step, samples were fixed in 250μL fixation buffer (BioLegend, 420801) for 20 minutes at 4°C. Flow cytometer experiments were performed with the MACSQuant16 (MACS Miltenyi Biotec). For compensation, the AbC Total Antibody Compensation Bead kit (Molecular Probes, A10497) was used upon incubation with both primary and secondary antibodies, following manufacturer recommendations. Data were analyzed with the FlowJo software (BD company).

### Statistical Analyses

For all the datasets, the nature of the distribution was tested with the D’Agostino & Pearson normality test. Pairwise comparisons of data following a normal distribution were carried out with Unpaired t-test; otherwise, the Mann Whitney U-test was performed. In order to assess the statistical significance of infection prevalence, Chi-square test was carried out. Statistical evaluation was carried out with GraphPad Prism v.8 (www.graphpad.com).

## Ethics Statement

All animal experimental procedures were reviewed and approved by the National Institute of Allergy and Infectious Diseases (NIAID) Animal Care and Use Committee under animal protocol LMVR4E. The NIAID DIR Animal Care and Use Program complies with the Guide for the Care and Use of Laboratory Animals and with the NIH Office of Animal Care and Use and Animal Research Advisory Committee guidelines. Detailed NIH Animal Research Guidelines can be accessed at https://oma1.od.nih.gov/manualchapters/intramural/3040-2/.

## Acknowledgment

We are thankful to Brian G. Bonilla and the other members of the sand fly insectary (LMVR/NIAID) for support. We are also thankful to Eva Iniguez and Ana Beatriz Barletta Ferreira (LMVR/NIAID) for scientific support. This research was supported by the Intramural Research Program of the NIH, National Institute of Allergy and Infectious Diseases. A.D. holds the Canada Research Chair on the Biology of Intracellular Parasitism. The funders had no role in study design, data collection and analysis, decision to publish, or preparation of the manuscript.

I.V.C.A. designed and performed the experiments and analyzed the data. J.O. supported flow cytometry experiments. D.C.W. performed parasite staining. T.R.W. helped with sand fly infections. C.M. performed sand fly insectary work. R.P.S. provided monoclonal antibody. A.D. and V.M.B provided the various *Leishmania* BH46 and BA262 lines. J.G.V. and S.K. were involved in the design, interpretation and supervision of this study. I.V.C.A wrote the first draft of the manuscript. J.G.V. and S.K. edited the manuscript.

---

**Figure S1.**
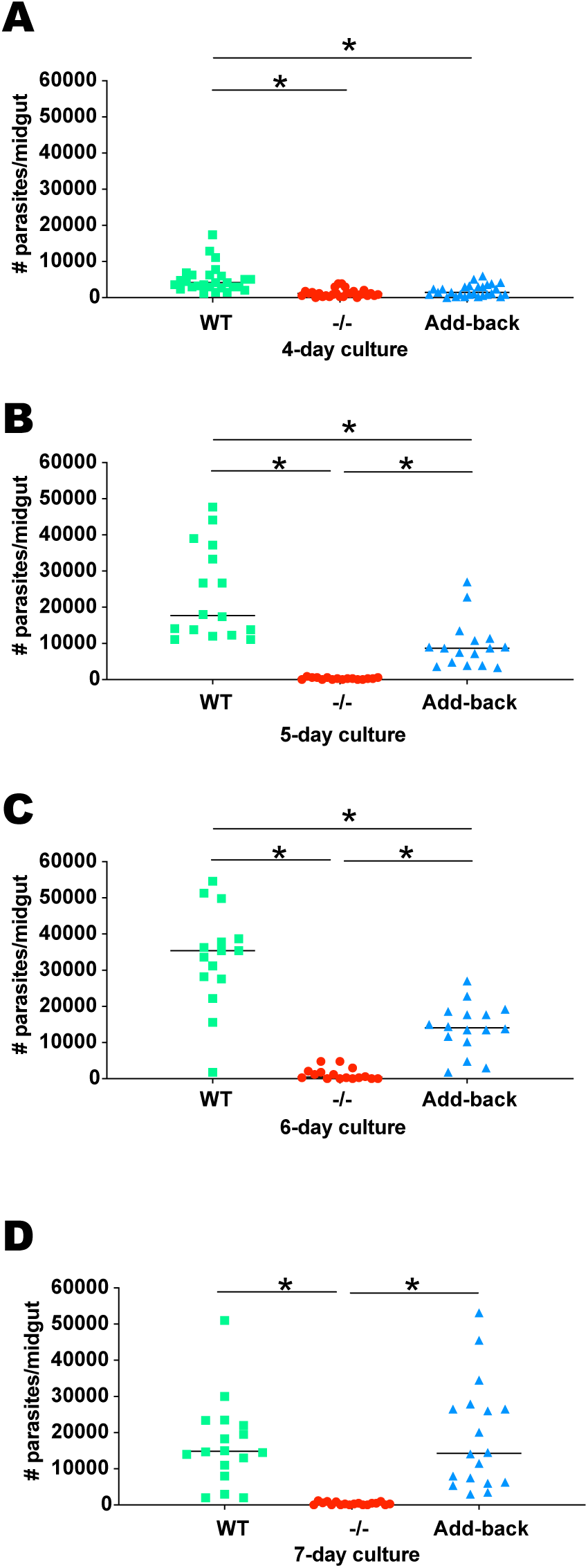
Binding of *Leishmania infantum* BH46 wild type, Δlpg1, and Δlpg1 + LPG1 lines to unfed *Lutzomyia longipalpis* midguts. (A-D). Cultures of *Le. infantum* BH46 WT (wild type), -/- (Δlpg1), and Add-back (Δlpg1 + LPG1) were harvested on days 4 (A), 5 (B), 6 (C), and 7 (D) and incubated with unfed midguts opened longitudinally along the anterior-posterior axis. n = 2. *, statistically significant at p < 0.05.

**Figure S2.**
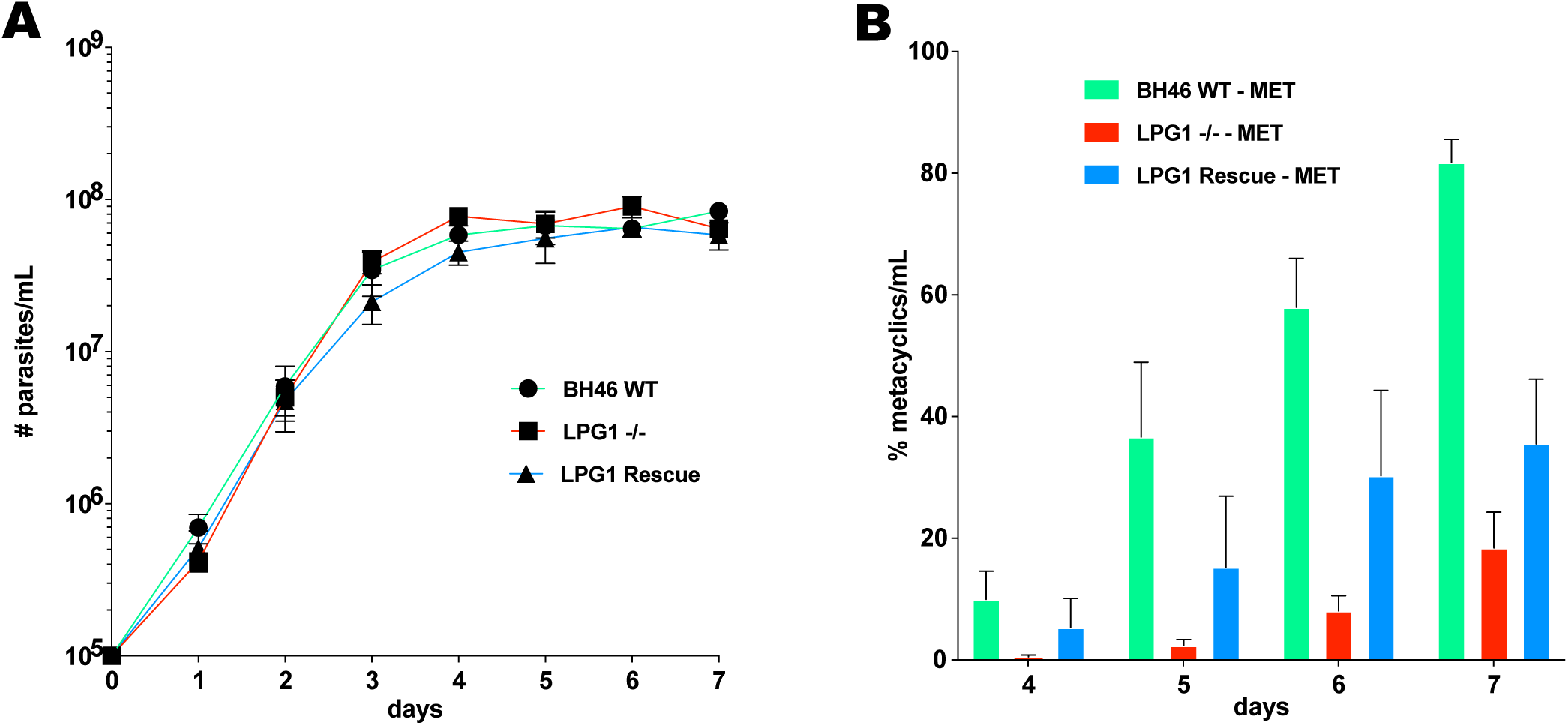
Growth curves of *Leishmania infantum* BH46 wild type, Δlpg1, and Δlpg1 + LPG1 parasites. (A) Growth curve *Le. infantum* BH46 strains in culture through day 7. n = 4. (B) Emergence of metacyclic *Le. infantum* BH46 strains in culture between days 4 and 10. n = 4.

**Figure S3.**
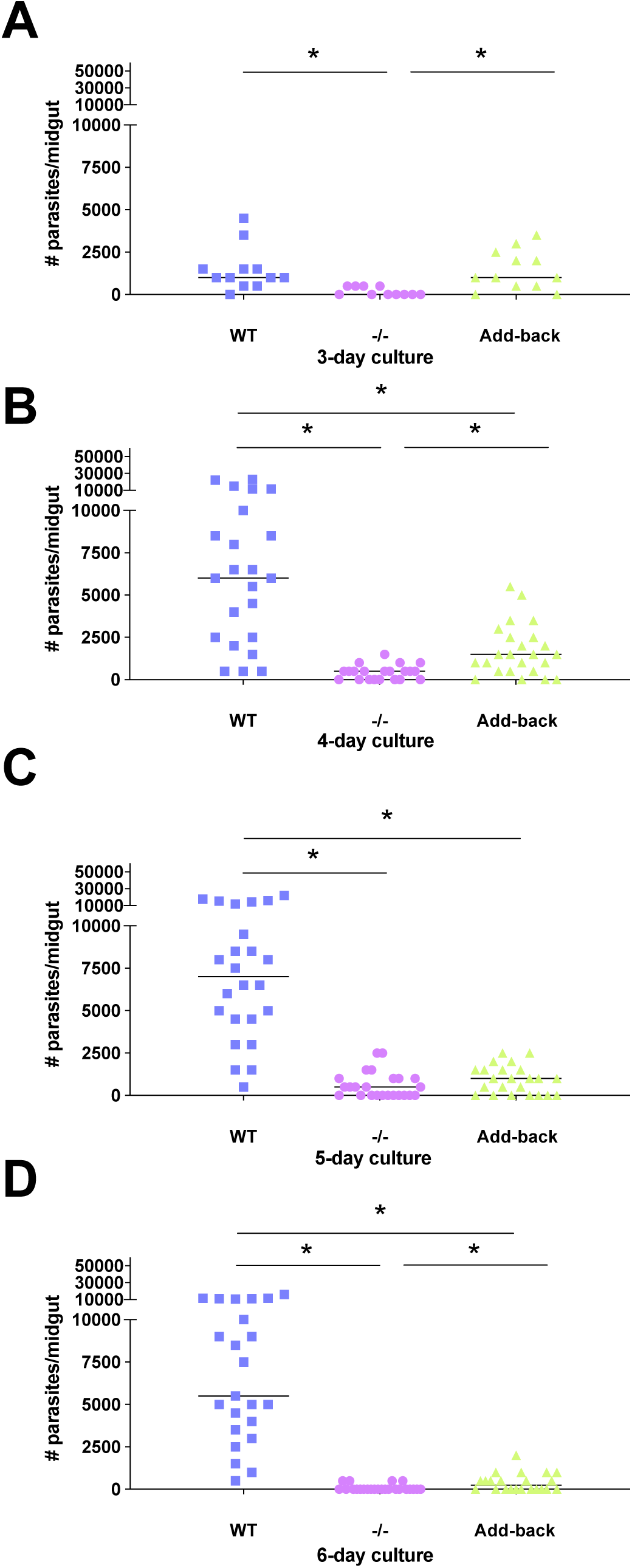
Binding of *Leishmania infantum* BA262 wild type, Δlpg1, and Δlpg1 + LPG1 lines to Lutzomyia longipalpis midguts. (A-D) Cultures of Le. infantum BA262 WT (wild type), -/- (Δlpg1), and Add-back (Δlpg1 + LPG1)were harvested on days 3 (A), 4 (B), 5 (C), and 6 (D) and incubated with unfed midguts opened longitudinally along the anterior-posterior axis. n = 2. *, statistically significant at p < 0.05.

**Figure S4.**
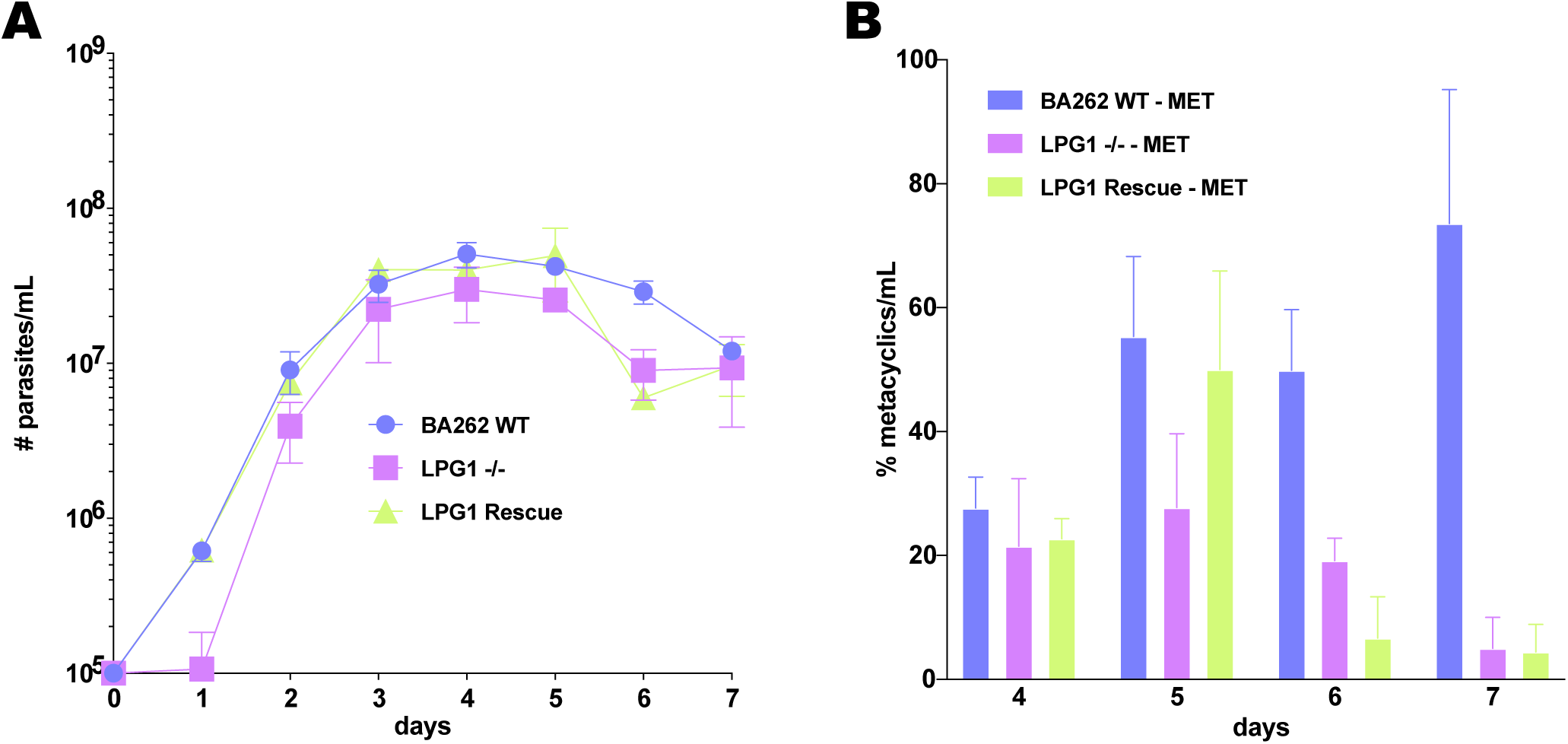
Growth curves of *Leishmania infantum* BA262 wild type, Δlpg1, and Δlpg1 + LPG1 parasites. (A) *Le. infantum* BA262 strains growth curve in culture through day 7 days. n = 3. (B) Emergence of metacyclics in cultures of *Le. infantum* BA262 strains between days 4 and 7. n = 3.

